# Embedding containerized workflows inside data science notebooks enhances reproducibility

**DOI:** 10.1101/309567

**Authors:** Jiaming Hu, Ling-Hong Hung, Ka Yee Yeung

**Author notes:** Correspondence should be addressed to K.Y.Y.

## Abstract

Data science notebooks, such as Jupyter, combine text documentation with dynamically editable and executable code and have become popular for sharing computational methods. We present *nbdocker*, an extension that integrates Docker software containers into Jupyter notebooks. *nbdocker* transforms notebooks into autonomous, self-contained, executable and reproducible modules that can document and disseminate complicated data science workflows containing code written in different languages and executables requiring different software environments.

---

With the generation of diverse and complex big data, computational method development and data analyses have become integral to research. Analytical protocols typically involve the execution of a series of computational tasks that are dependent on code, parameters, environment, installation and setup, that are not easily described by text inside a traditional laboratory notebook or in a published “Materials and Methods” section. Data science notebooks such as Jupyter notebooks offer a partial solution as they allow for inclusion of executable live code and documentation. Each Jupyter notebook is a web application that is divided into markdown (text) and code cells that can be modified and run independently inside the notebook ^1,2^. All modifiable code cells in a notebook must be in the same language but Jupyter has kernels supporting over 100 programming languages including R, Python, Ruby, Javascript, C++ and Perl ^1,3,4^. The integration of editable code with the scientific rationale and narrative facilitates the documentation, dissemination and adoption of computational methodologies. As a result, Jupyter notebooks have become extremely popular, with over 1.7 million Jupyter notebooks shared on the public GitHub code repository ^4^ covering a wide variety of scientific disciplines. These interactive notebooks have become particularly useful in bioinformatics for documenting complicated workflows and sharing analytical protocols between collaborators with different backgrounds ^5-7^. Many bioinformatics software tools such as GenePattern ^8^ and Galaxy ^5^ have built-in support for Jupyter notebooks.

A major drawback of Jupyter notebooks is that they are not autonomous. Although the code in code cells is modifiable and executable, execution often requires the installation of additional software, libraries, frameworks and packages by the user. For bioinformatics workflows, there are usually many components or modules, each of which executing a different tool that requires potentially different computing environment and software dependencies. One approach to this problem is to use software containers such as Docker containers to encapsulate each computing environment. Docker containers wrap the executables and scripts inside a custom software environment, avoiding conflicts between different components and thus, eliminating the need for users to install and manage all the software dependencies. Dockerized components are completely isolated and modular and will yield identical results regardless of the platform. When a Docker command is run, Docker will search for and automatically download the specified container from its public repository, eliminating the need for installation of diverse and possibly conflicting software. As a result, Docker container technology is rapidly gaining popularity in biomedical research and has been used to enhance both the portability and reproducibility of complicated workflows^9^.

Another limitation of Jupyter notebooks is that each notebook is limited to one kernel supporting a single programming language. This is a problem since bioinformatics workflows typically consist of a series of tasks that are written in different programming languages. There is some limited support for multiple languages with a single notebook. rpy2 embeds an R interface inside a Python process ^10^. Beaker notebooks (currently “beakerX”) can also support multiple languages but requires that the user to create a REST server to wrap the application ^11^.

Another possible workaround is to create a custom kernel that provides the software requirements for all the components. Users would still have to choose a single language and execute other commands using mechanisms provided by that language (e.g. the subprocess module of Python). However, the portability and modularity of the workflow are limited by the use of a custom kernel.

Using Docker containers inside Jupyter notebooks offers a more robust and flexible approach. Currently, this is possible using Jupyter’s ability to embed shell commands within code cells. However the syntax and availability of these commands varies between different kernels^12^. For example, preceding a shell command with an exclamation point “!” works in the Python kernel but not in the R kernel. Here, we present “nbdocker”, a Python/Javascript extension to Jupyter notebooks that allows for different Docker containers to be executed inside Jupyter notebooks in the same manner regardless of the kernel used. nbdocker is an extension that integrates a Docker management user interface (UI) into Jupyter. The user can embed a set of Docker commands as clickable buttons inside markdown (text) cells. Specifically, “nbdocker” provides a point-and-click Docker management UI to pull a Docker image from a registry such as DockerHub or a local image, keep a record of running Docker containers, document and execute a Docker container in the history. The user can also check the status of running containers with a single click.

We illustrate the utility of “nbdocker” using an established RNA sequencing (RNA-seq) data processing workflow using kallisto^13^ and sleuth ^14^ In this workflow ^15^, the first step is to download data files from NCBI’s Short Read Archive (SRA) database using a Python script that calls the NCBI SRA toolkit. Next, the reference human transcriptome is downloaded and indices are generated using the annotation file. The reads are then aligned to the reference and the abundance of the transcripts are quantitated using kallisto, a binary executable compiled from C++ source code. Finally, differentially expressed genes under different experimental conditions are identified Sleuth ^14^ which is written in R. To replicate this workflow from the original published methods description requires the configuration of multiple computing environment and installation of multiple software tools, including kallisto, sleuth, python, R, SRA tools and Bioconductor packages. Most importantly, the versions of all software must be compatible with each other and with the reader’s hardware and operating system. Without “nbdocker”, the user will have to manually install and execute selected steps in this workflow outside the Jupyter notebook. One possibility is to execute the data download and kallisto pseudoalignment steps outside the notebook and represent the differential expression step in a notebook running the R kernel. In contrast, “nbdocker” allows each module to be represented as an independent Docker container, and hence, multiple programming languages and conflicting dependencies can be encapsulated within a single notebook. Most importantly, the history of Docker commands are all contained in the “nbdocker” notebook such that simply sharing the notebook file (.pynb file) will allow a collaborator to reproducibly execute and modify the workflow without any additional download or installation. The differences between these different methods of documenting computational workflows are shown in **Figure 1**.

In this paper, we present a Jupyter extension called “nbdocker” that brings the modularity, portability and reproducibility of software containers to Jupyter notebooks (see **Table 1** for a summary of features). Complex workflows with multiple modules and different software requirements can be encapsulated within a single nbdocker notebook. In addition, “nbdocker” offers an interactive computing environment that allows easy sharing, modification and point-and-click reproducible execution of documented Docker commands, thus facilitating the communication of complicated data science protocols and collaboration between diverse groups of researchers.

**Figure 1.**
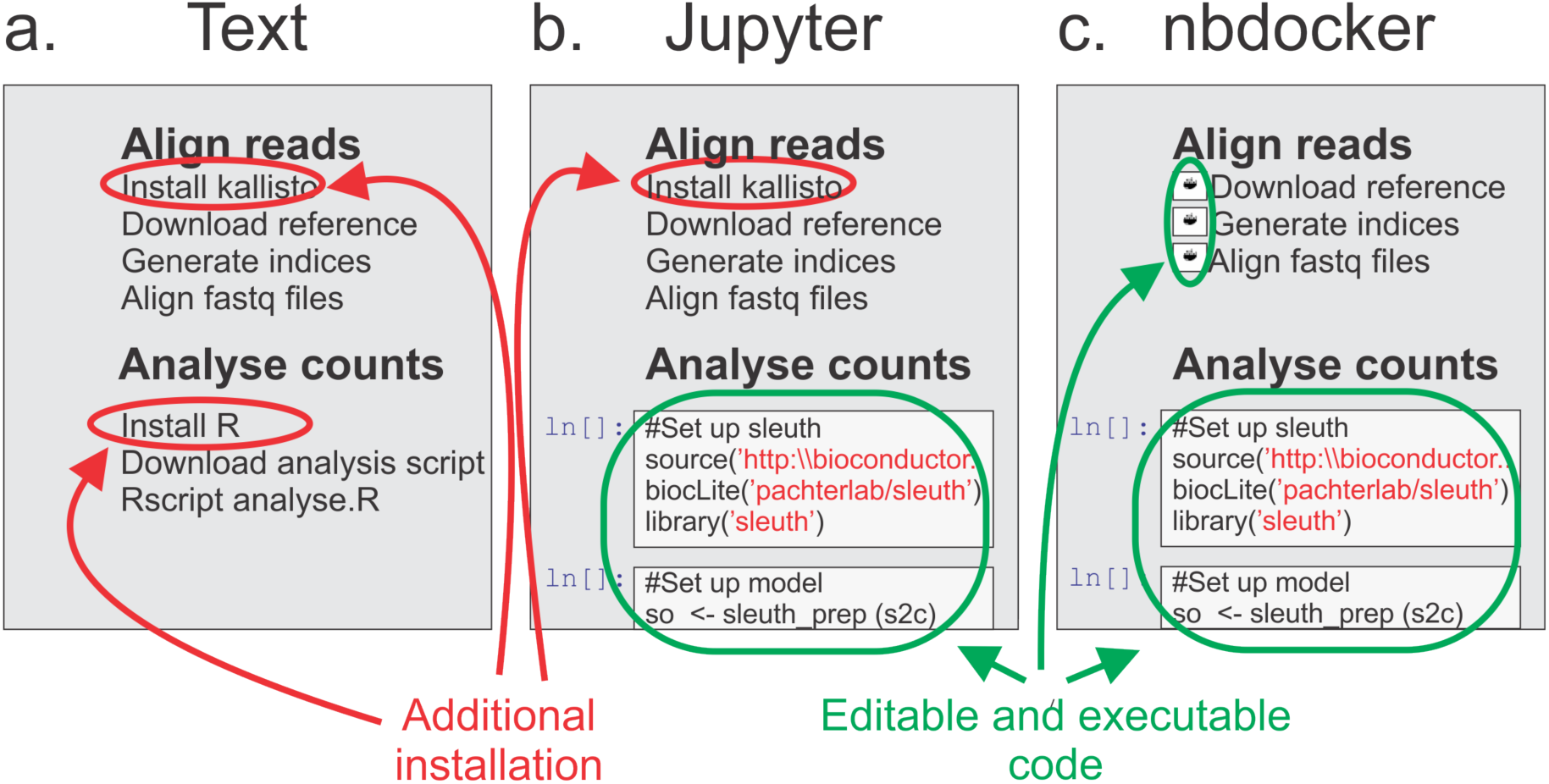
Complex bioinformatics workflows illustrate differences between static text documentation, Jupyter notebook and Jupyter notebook with nbdocker extension. A RNA-seq processing workflow workflow using kallisto for pseudoalignment and sleuth for identification of differentially expressed genes is used here as an example bioinformatics workflow.

a. Using static text description, users must configure the computing environment and install all software tools (circled in red). Since the code in the text file is not executable, users may unintentionally omit parameters or instructions that match the latest version of the code.
b. Jupyter notebook contains embedded executable code cells (circled in green). This ensures that the code that was actually run is accurately represented in the notebook. The code in the notebook can be modified facilitating customization. However, the Jupyter notebook is limited to a single kernel that only supports the R (sleuth) part of the pipeline. The user must install and execute the data download and kallisto steps outside the notebook.
c. Finally, “nbdocker” adds the ability to embed Docker commands as clickable buttons in markdown cells. Users can execute the Docker commands corresponding to the pseudoalignment (kallisto) task with a point-and-click user interface. The Docker commands can be documented and modified. Using “nbdocker”, the entire workflow is modularized and contained within a single notebook that can be shared, modified and reproducibly executed without installation of any additional software.

**Table 1.**
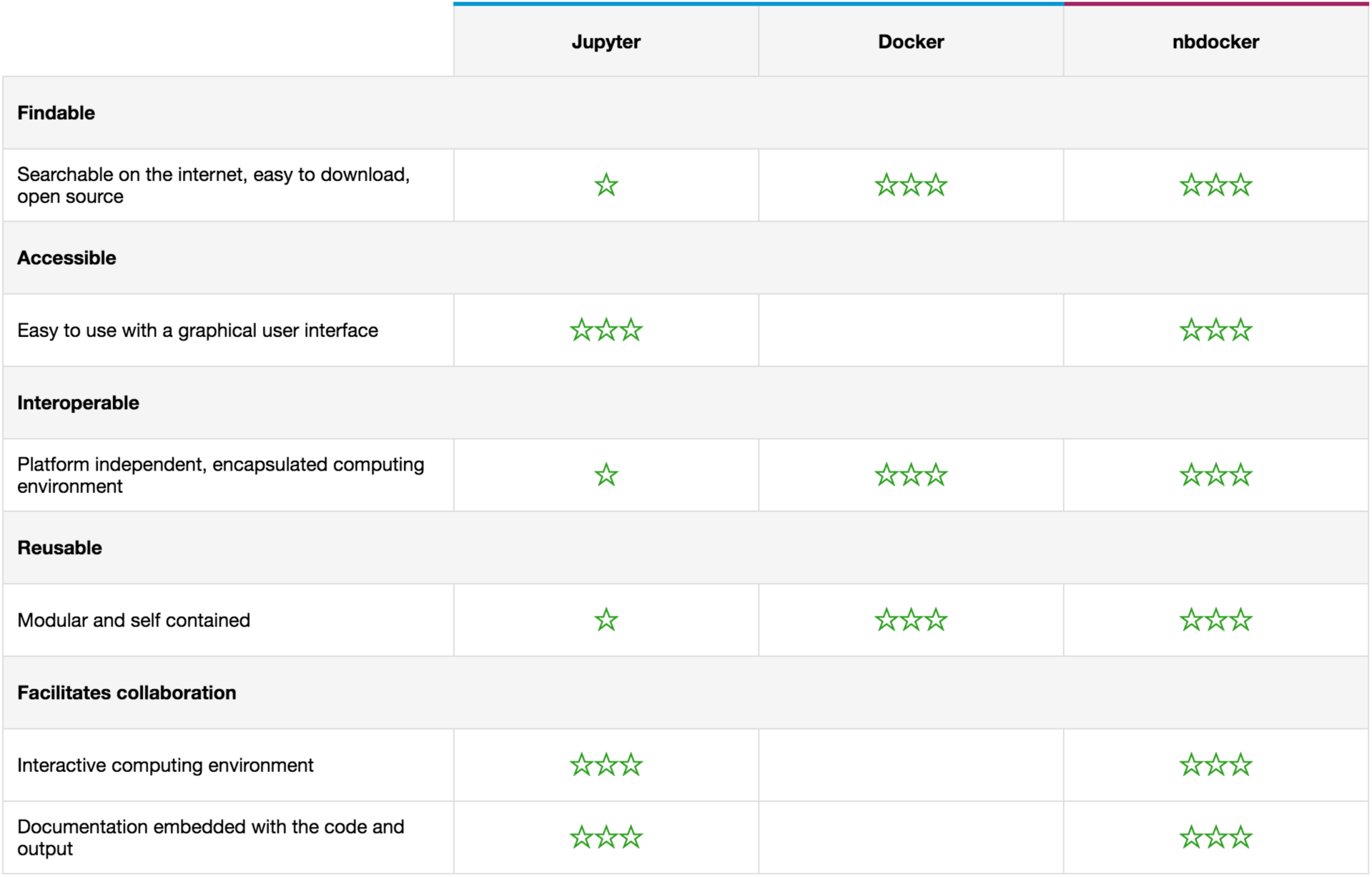
Comparison of features of Jupyter, Docker and nbdocker. Three stars mean that the requirement is strongly satisfied. One star means that the requirement is weakly satisfied.

|  | Jupyter | Docker | nbdocker |
| --- | --- | --- | --- |
| **Findable** |  |  |  |
| Searchable on the internet, easy to download, open source | ☆ | ☆☆☆ | ☆☆☆ |
| **Accessible** |  |  |  |
| Easy to use with a graphical user interface | ☆☆☆ |  | ☆☆☆ |
| **Interoperable** |  |  |  |
| Platform independent, encapsulated computing environment | ☆ | ☆☆☆ | ☆☆☆ |
| **Reusable** |  |  |  |
| Modular and self contained | ☆ | ☆☆☆ | ☆☆☆ |
| **Facilitates collaboration** |  |  |  |
| Interactive computing environment | ☆☆☆ |  | ☆☆☆ |
| Documentation embedded with the code and output | ☆☆☆ |  | ☆☆☆ |

## Methods

### Implementation details

“nbdocker” has two components: a Jupyter server extension (back-end) and a Jupyter notebook extension (front-end). The server extension reserves three URL’s : */docker, /dockerpull* and */dockerbuild. /docker*is used to run basic docker commands (e.g. list images/containers and pull/build image submission) while the URL /dockerpull and /dockerbuild are reserved for building/pulling docker image. The notebook front-end is implemented in javascript (JS), renders the JSON data provided by the back-end and interacts with users. **Figure S1** shows the overall architecture of “nbdocker”.

### nbdocker Engine: server extension

The server extension uses the Docker API for python (docker-py) ^17^ to interact with the Docker engine. Docker commands are executed through the *unix:///var/run/docker.sock* socket on linux/unix and *npipe:////./pipe/docker_engine* socket on Windows. Some jobs such as pulling or building a Docker image can be quite lengthy. We implemented a session manager to run these jobs as threads and return control to the user. The session manager is designed to be used by multiple notebooks. A separate uuid is provided for each job allowing them to be tracked individually. The session will terminate when all jobs have finished. **Figure S2** summarizes the process of building a Docker image in “nbdocker”.

### Docker management UI: notebook extension

The front end is based on notebook’s JS library that is used to interact with users. We added two additional libraries: xterm.js ^18^ which renders image building logs and progressbar.js ^19^ which shows the progress of lengthy jobs such as pulling images. AJAX is used for one-step commands such as listing images. This prevents the webpage from refreshing. For longer jobs such as pulling or building an image, a POST mechanism is used to pass the Docker image or Dockerfile to the server extension. We then monitor the server extension for events that we pass to the EventSource object to display the progress to the user.

To integrate nbdocker inside the Jupyter markdown cells we implement a custom rendering function for markdown cells. The custom function searches for the keyword {nbdocker#<history_id>} in the cells and allows other content pass through to the functions that are responsible for rendering the markdown cells. Upon finding the keyword, it is rendered as whale shaped badge. Both the <cell_id> and <history_id> are bound to the click event of this badge allowing us to identify which badge has been clicked by the user and which set of Docker commands should be run in response.

### Docker run history

nbdocker records the directory mapping, port mapping and docker commands when the user runs a docker container through nbdocker. These histories are saved along with the Jupyter notebook file (.ipynb). The front-end notebook extension also sends the current working notebook name to the server where the global history dictionary was maintained. The appropriate histories are then written in JSON format to the matching ipyb file. nbdocker monitors clicks on badges in the markdown cells. When a replay event is triggered by a badge click, a request is sent to the server to retrieve the matching history. The Docker container id will be written into the metadata of the markdown cell where the badge was clicked. A status bar is attached to the bottom of the current cells to indicate the running status of the container(s) launched.

### Case study: RNA-seq data processing workflow using STAR and edgeR

We also include a second case study on another RNA-seq work documented in Bioconductor ^16^. Specifically, the Love *et al.* workflow used STAR to align short reads to the reference genome and DESeq2 to infer differentially expressed genes. STAR ^17^ is written in C++ while DESeq2 ^18^ is a R package in Bioconductor. The existing workflow documented in Bioconductor provides all the R code to perform the differential expression tasks after the BAM files are generated from the alignment step. **Figure S3** illustrates this case study.

### Software availability

The source code is publicly available on GitHub at https://github.com/BioDepot/nbdocker. A pre-built Docker image is publicly available on DockerHub with “nbdocker” extension installed: https://hub.docker.com/r/biodepot/nbdocker/

**Figure S1.**
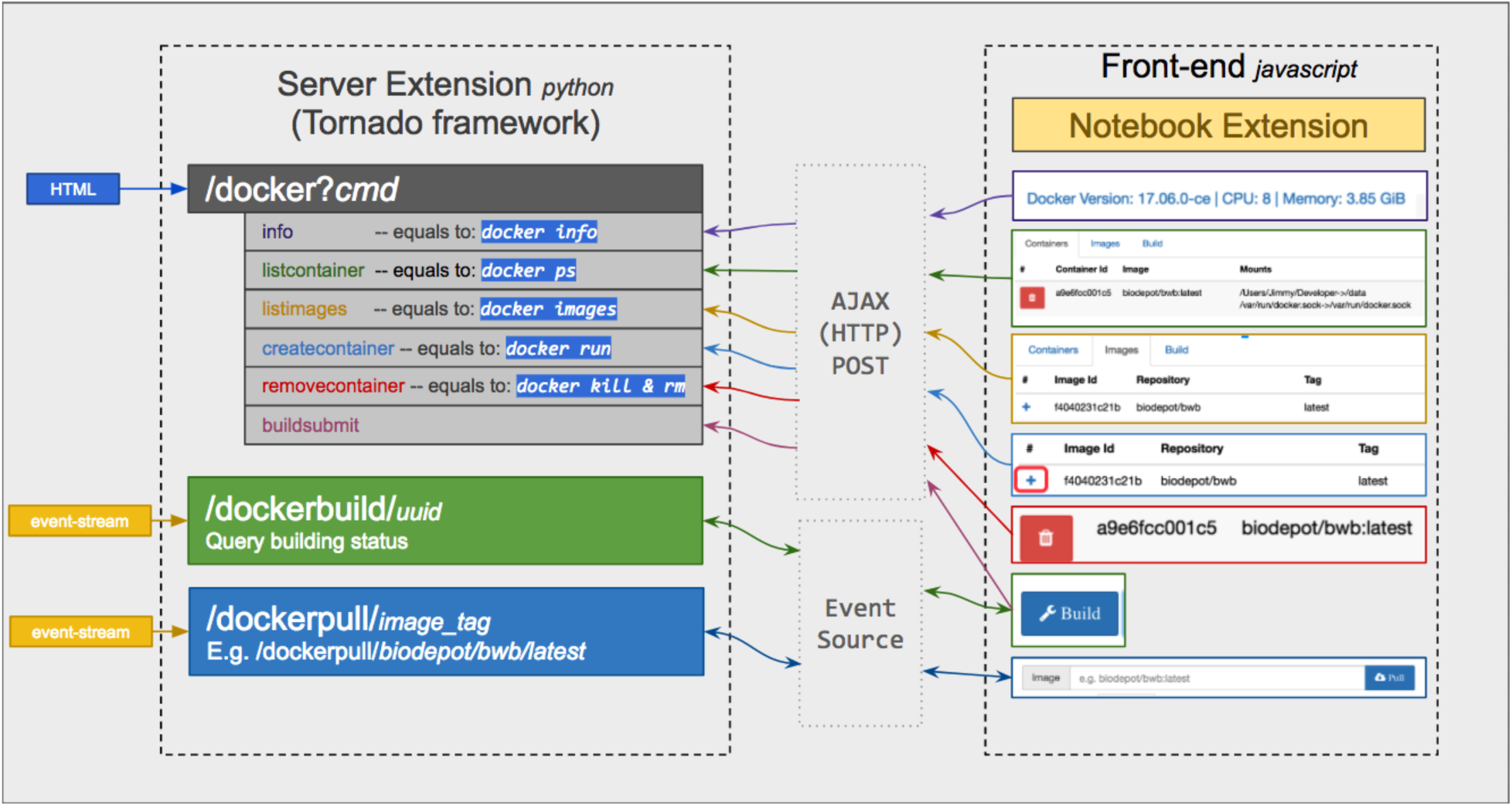
Software architecture of “nbdocker”.

**Figure S2.**
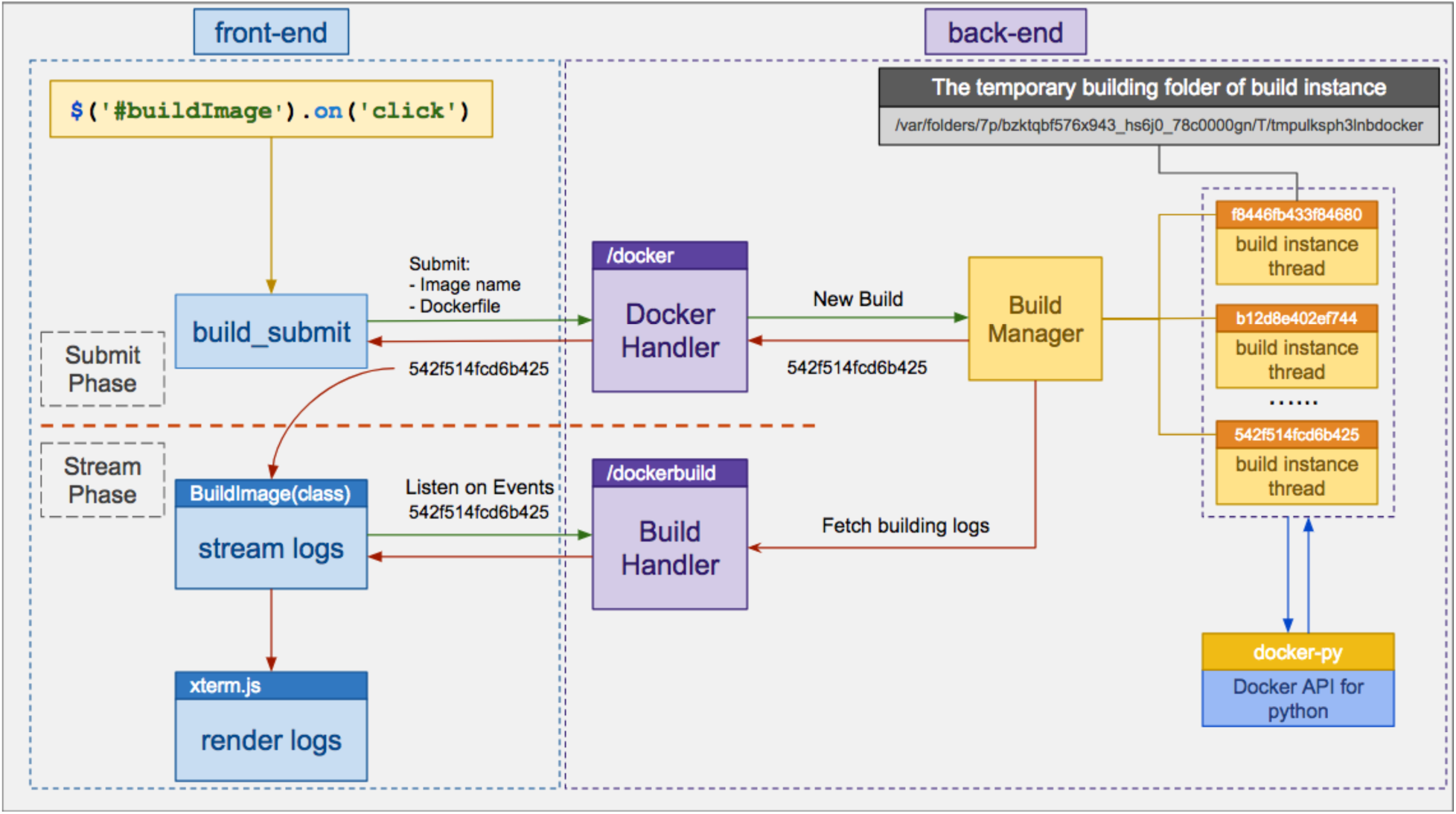
The process and events of building a Docker image in “nbdocker”.

**Figure S3.**
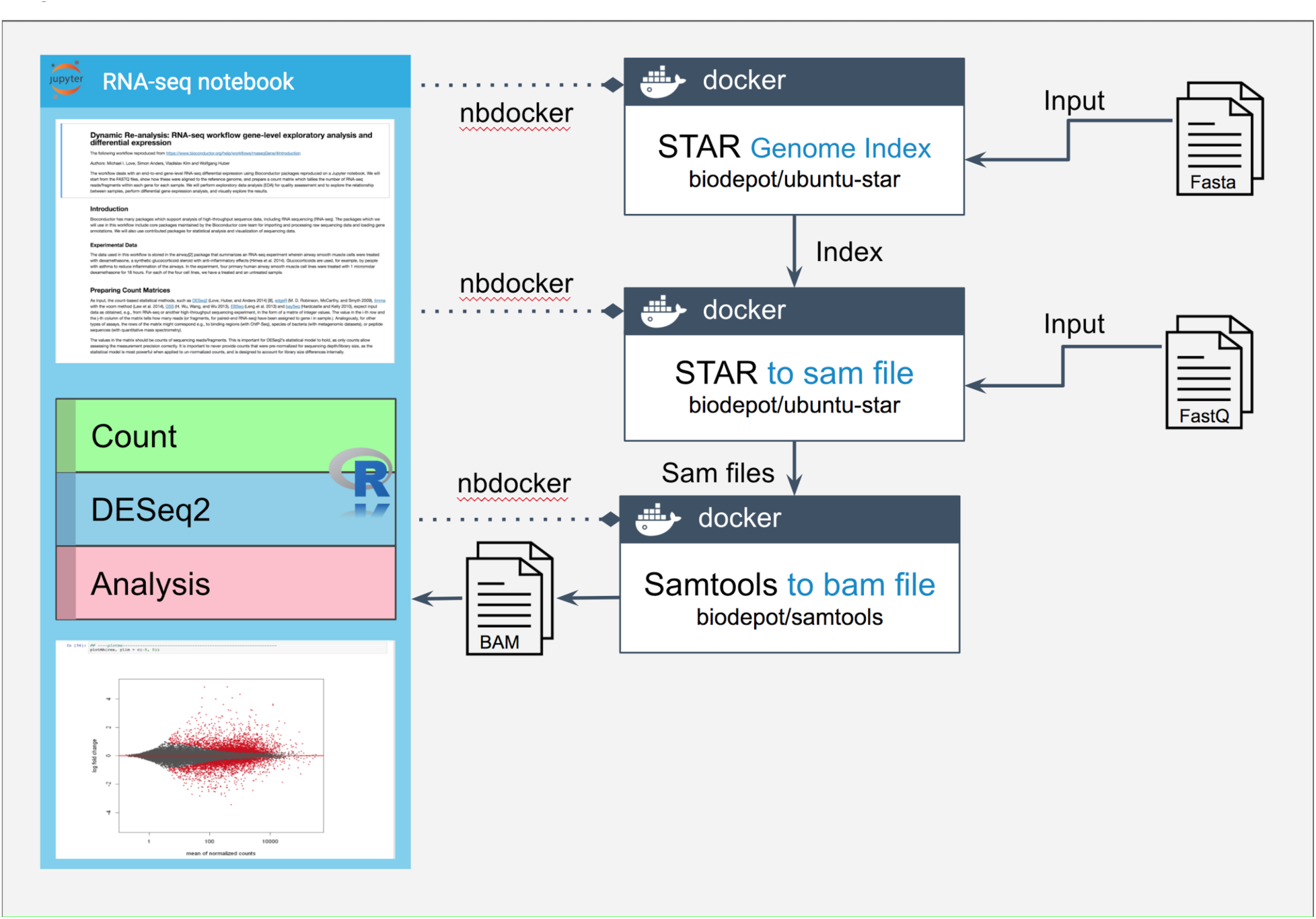
An additional case study in which RNA-seq data are processed using STAR and edgeR.

## Acknowledgements

We would like to thank Dr. Wes Lloyd for helpful discussions in group meetings. We would like to thank Mr. Fang Chen for researching the different Jupyter magic commands and working on earlier implementations of the saving docker histories. L.H.H. and K.Y.Y. are supported by NIH grants U54HL127624 and R01GM126019. J.H. is supported by U54HL127624. We would also like to thank the Center for Data Science and the Institute of Technology at University of Washington Tacoma for the purchase of a computer server.

## Author Contributions

J.H. designed, implemented and tested the Jupyter extension “nbdocker”. J.H, L.H.H and K.Y.Y created the Jupyter notebooks for the RNA-seq case study. L.H.H. and K.Y.Y. drafted the manuscript. L.H.H and K.Y created the tables and figures in the manuscript. J.H. created the figures in Online Methods, wrote user documentation and created tutorial videos. L.H.H. tested and refined the containers. K.Y.Y. and L.H.H. coordinated the study. All authors edited the manuscript.

## References

1. Pérez, F. & Granger, B.E. IPython: A System for Interactive Scientific Computing. Computing in Science and Engineering 9, 21–29 (2007).

2. Kluyver, T. et al. in Positioning and Power in Academic Publishing: Players, Agents and Agendas. (eds. F. Loizides & B. Schmidt) 87–90 (2016).

3. Jupyter kernels. https://github.com/jupyter/jupyter/wiki/Jupyter-kernels

4. JupyterLab is Ready for Users. https://blog.jupyter.org/jupyterlab-is-ready-for-users-5a6f039b8906

5. Gruning, B.A. et al. Jupyter and Galaxy: Easing entry barriers into complex data analyses for biomedical researchers. PLoS computational biology 13, e1005425 (2017).

6. Jupyter Genomics: A collection of Jupyter notebooks authored by the UCSD Center for Computational Biology & Bioinformatics https://github.com/ucsd-ccbb/jupyter-genomics

7. Wang, Z. & Ma’ayan, A. An open RNA-Seq data analysis pipeline tutorial with an example of reprocessing data from a recent Zika virus study. F1000Research 5, 1574 (2016).

8. Reich, M. et al. The GenePattern Notebook Environment. Cell systems 5, 149–151 e141 (2017).

9. Silver, A. Software simplified: Containerization technology takes the hassle out of setting up software and can boost the reproducibility of data-driven research. Nature 546, 173–174 (2017).

10. rpy2. https://rpy2.bitbucket.io/

11. Beaker. http://beakernotebook.com/

12. IPython: built-in magic commands. http://ipython.readthedocs.io/en/stable/interactive/magics.html-line-magics

13. Bray, N.L., Pimentel, H., Melsted, P. & Pachter, L. Near-optimal probabilistic RNA-seq quantification. Nature biotechnology 34, 525–527 (2016).

14. Trapnell, C. et al. Differential analysis of gene regulation at transcript resolution with RNA-seq. Nature biotechnology 31, 46–53 (2013).

15. kallisto and sleuth walkthrough. https://github.com/pimentel/bears_iplant/blob/master/README.md

16. Love, M.I., Anders, S., Kim, V. & Huber, W. RNA-Seq workflow: gene-level exploratory analysis and differential expression. F1000Research 4, 1070 (2015).

17. Dobin, A. et al. STAR: ultrafast universal RNA-seq aligner. Bioinformatics 29, 15–21 (2013).

18. Love, M.I., Huber, W. & Anders, S. Moderated estimation of fold change and dispersion for RNA-seq data with DESeq2. Genome biology 15, 550 (2014).

